# Assessing impacts of mitochondrial dysfunction on axonal microtubule bundles as potential mechanism in neurodegeneration

**DOI:** 10.1101/2025.05.11.653329

**Authors:** Scott Murray-Cors, Milli Owens, Yu-Ting Liew, Maureece Day, William Cairns, Andreas Prokop

**Author notes:** shared first authorship (Scott Murray-Cors, Milli Owens).

## Abstract

Mitochondrial dysfunction is an important cause for neurodegeneration, often associated with dyshomeostasis of reactive oxygen species, i.e. oxidative stress. However, apart from ATP production, mitochondria have many other functions the aberration of which may impact neurons in very different ways. Oxidative stress can cause the deterioration of axonal microtubule bundles, thus critically affecting the highways for life-sustaining transport and providing a potential path to neurodegeneration. We recently found that aberrant transport of mitochondria can have this effect by causing oxidative stress. We therefore asked which aberrations of mitochondrial physiology might impact microtubules, which of these might explain the observed consequences of aberrant mitochondrial transport, and whether mitochondria-induced microtubule phenotypes are always mediated by oxidative stress. Using one consistent *Drosophila* primary neuron system, we deleted 13 different mitochondrial factors known to be detrimental for neurons *in vivo*. Losses of five factors caused MT damage, all involving oxidative stress, hence supporting the path from mitochondria via oxidative stress to microtubule deterioration; we discuss Sod2 as potential candidate explaining effects of mitochondrial transport aberration. However, the loss of eight factors - seven of them important mitochondrial morphogenesis regulators - caused no microtubule damage, suggesting potential oxidative stress-independent pathways.

**Summary Statement:** Assessing mutant effects of 13 mitochondrial factors on axonal microtubule organisation to unravel potential mechanisms underpinning neurodegeneration

## Introduction

Neurodegenerative disorders are an important socioeconomic challenge to modern ageing societies (GBD 2016 Neurology Collaborators, 2019). One major cellular cause often highlighted in this context is the dysregulation of mitochondria (Wang et al., 2021), especially the excessive generation or inappropriate release of harmful reactive oxygen species (ROS) as a side product of the electron transfer chain (ETC) that drives oxidative phosphorylation (OXPHOS; Fig.1; Andreyev et al., 2005; Murphy, 2008; Zorov et al., 2014).

**Fig. 1.**
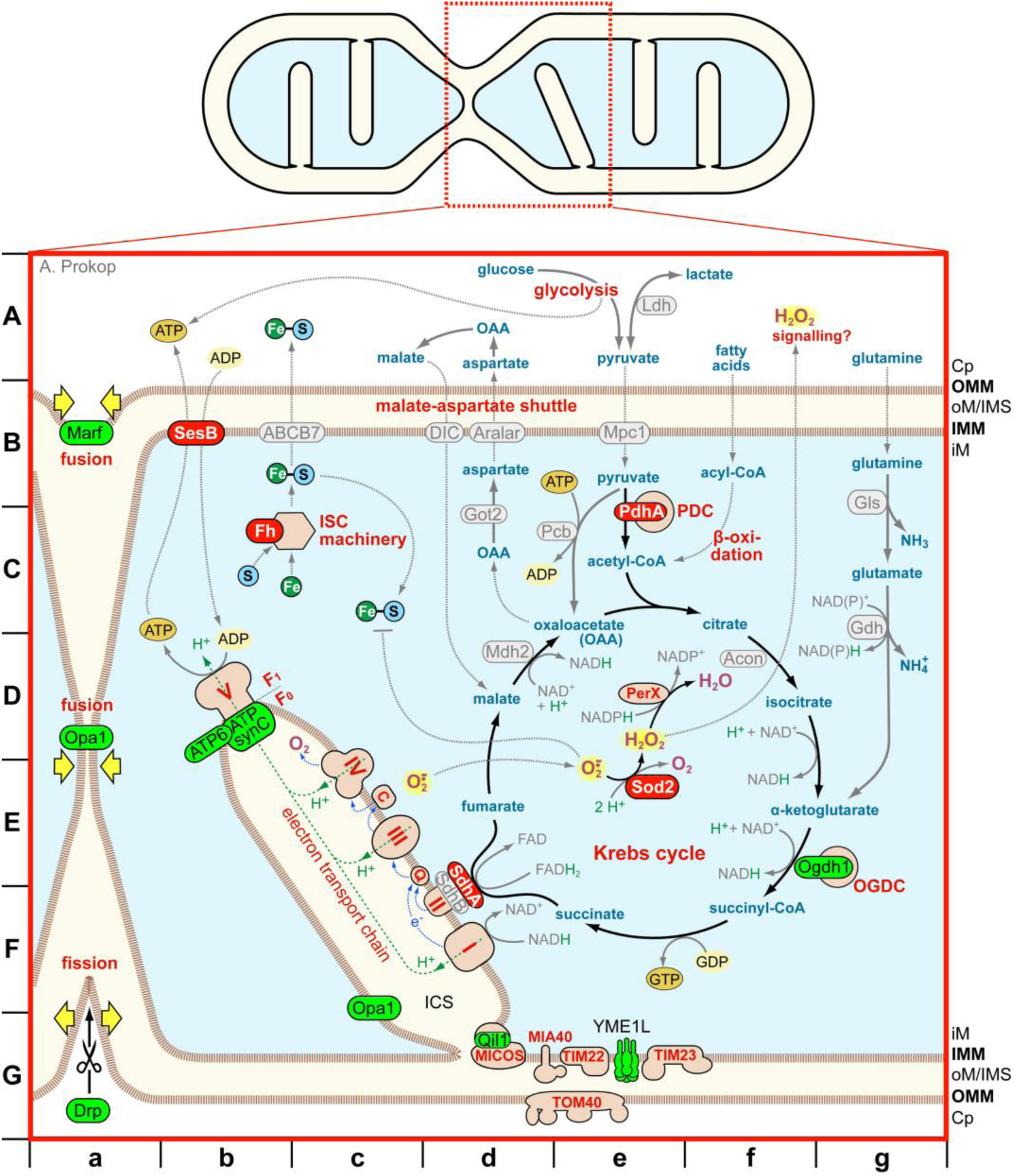
Processes and components referred to in the text that mediate mitochondrial physiology and dynamics. The sketch on top represents a mitochondrion during fusion/fission, with the red box indicating the position of the close-up shown below. Different compartments and membranes are colour-coded: cytoplasm (Cp) in white, outer mitochondrial matrix/inter-membrane space (oM/IMS) and inter-cisternal space (ICS) in light beige, inner mitochondrial matrix in light blue, inner and outer mitochondrial membranes (IMM, OMM) as stippled brown lines; for further acronyms see the dedicated abbreviation list. Protein complexes are shown in pink with red outlines; identified proteins are either in light green/dark red with black outline (proteins manipulated in this study without/with MT-curling), or in light-grey with grey outline (mentioned in the text but not investigated here). Organic and inorganic components are colour coded as follows: organic metabolites in blue, protons in green, oxygen derivatives in magenta (those being ROS highlighted in yellow), co-enzymes of redox reactions in grey, iron (Fe) in dark-green circles, sulphur (S) in blue circles, ATP/GTP in dark orange circles, ADP/GDP in light orange circles. Chemical reactions are shown as solid black or grey arrows, spatial translocations as stippled grey arrows. Letters on the left and at the bottom outside the box are grid coordinates referred to in the text (chevron followed by italics letters).

However, mitochondria are far more than the cell’s powerhouse. For example, astrocytes, activated immune cells, tumour cells or cells during migration often circumvent the resource-efficient but slow process of oxidative phosphorylation; they instead switch to aerobic glycolysis as resource-hungry but fast and locally available means of ATP generation, which also churns out pyruvate molecules as metabolic building blocks or turning them into lactate as secondary energy source (see *A/de* grid in Fig.1; Bhattacharya et al., 2020; Krawczyk et al., 2010; Magistretti and Allaman, 2018; Rodríguez-Prados et al., 2010). It was even reported that mitochondria in neuronal axons often act as ATP sink rather than source (Hirabayashi et al., 2024). Other roles of mitochondria are coming to light (Pfanner et al., 2019). For example, they are essential to produce iron-sulphur clusters as obligatory components of many enzymes in mitochondria, the cytoplasm or nucleus (Marelja et al., 2018; Shi et al., 2021). Mitochondria are discussed as calcium-buffering stores at synapses (Walters and Usachev, 2023), play key roles in programmed cell death involving the release of signals such as cytochrome c or mitochondrial DNA (Bonora et al., 2022; Glover et al., 2024; Khatun et al., 2024) and display direct contacts with other organelles or with each other as means of cross-regulation (Picard et al., 2015; Voeltz et al., 2024). Furthermore, mitochondria appear to respond to oxidative and metabolic states or requirements of cells by adapting their activities, changing shape, sending signals to the cytoplasm, positioning themselves into subcellular locations as extreme as the tips of filopodia or even transiting between cells (Fig.1>*A/f*); in this way they seem to help maintain cellular homeostasis or change the activity states of cells (Borcherding and Brestoff, 2023; D’Angelo et al., 2023; Marlar-Pavey et al., 2025; Palma et al., 2020; Sturm et al., 2024).

To add to this complex picture, we recently observed in *Drosophila* primary neurons that axonal transport deficits of mitochondria lead to ROS dyshomeostasis which, in turn, triggers deterioration of axonal microtubule (MT) bundles displaying as severe curling and criss-crossing of individual MTs (Fig.2). Notably, this phenotype occurred in axons even when mitochondria remained far away in cell bodies (Liew et al., 2025). Therefore, the wrong positioning or even absence of mitochondria seems to have a negative impact on axonal MT bundles.

**Fig. 2.**
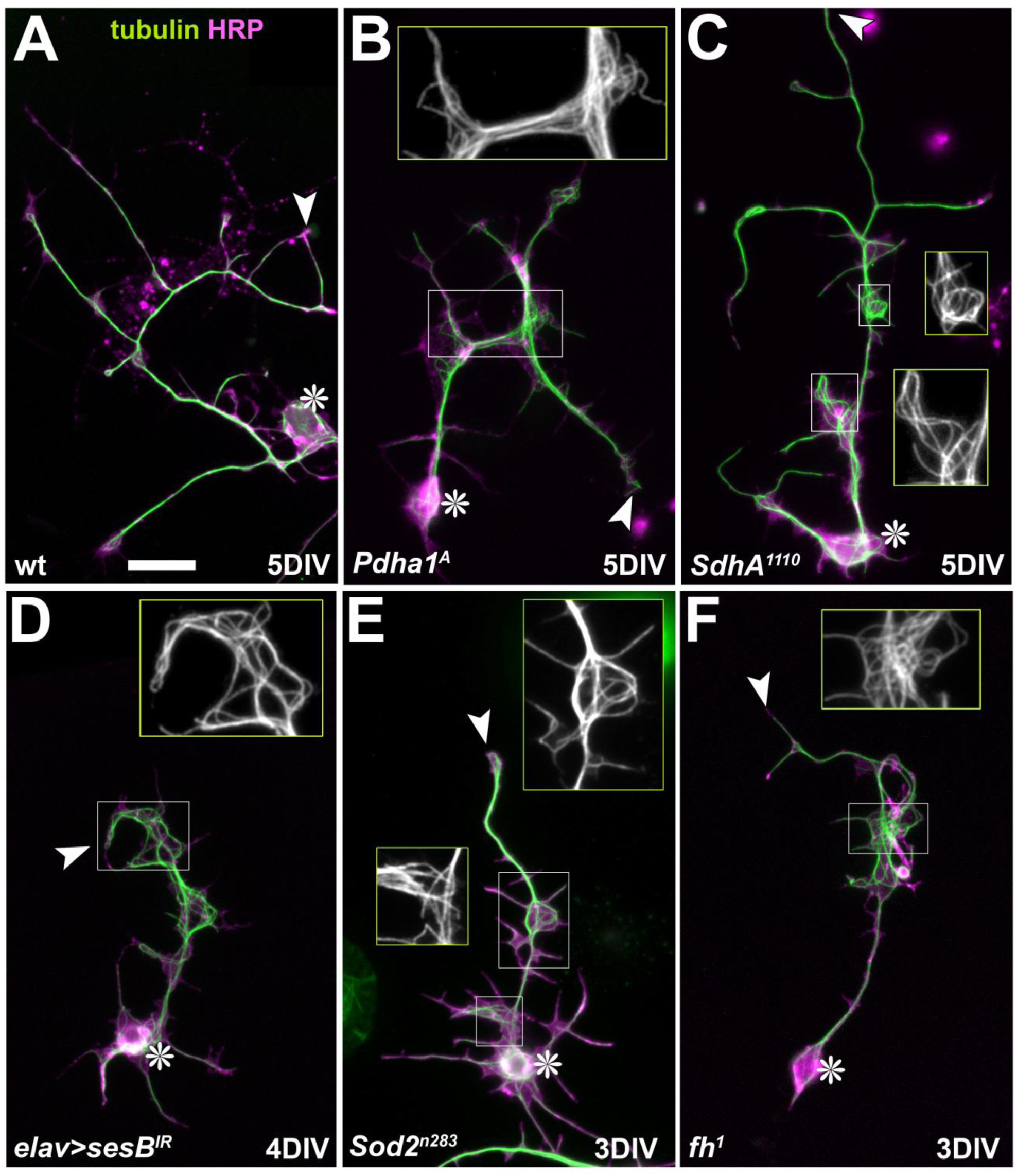
Representative images of MT-curling observed in mutant neurons. *Drosophila* primary neurons of different genotypes (indicated bottom left; see text) cultured for 3, 4 or 5 days *in vitro* (DIV) stained for tubulin (green) and the neuronal surface marker HRP (magenta). Cell bodies are indicated by asterisks, axon tips by arrow heads, and emboxed areas are shown as two-fold magnified insets (tubulin channel only). Scale bar in A represents 20µm in all images.

These findings are relevant because they establish a link from mitochondrial function to MT bundles as the essential lifeline of axons. MT bundles run uninterrupted from the neuronal cell bodies to the tips of axons and serve as the highways for cargo transport required to sustain axonal cell biology; when interrupting these bundles, they become an axon’s Achilles heel on path to degeneration (Okenve-Ramos et al., 2024; Prokop, 2021; Smith et al., 2023). We therefore wondered (1) what malfunctions of the complex mitochondrial biology can lead to MT bundle damage, (2) whether this is always mediated by harmful ROS, and (3) how the absence of mitochondria from axons might lead to MT curling.

To address these questions, we carried out a pilot study systematically assessing the functional loss of 13 well-conserved mitochondrial proteins (Tab.1; Brischigliaro et al., 2023; Marelja et al., 2018). All studies were carried out in one consistent *Drosophila* primary neuron system and using MT-curling as the key readout (Hahn et al., 2021; Prokop et al., 2013). Although all conditions are known to be debilitating for neurons *in vivo*, eight genes could not be linked to MT curling in cultured neurons, with seven of them being important morphogenesis regulators of mitochondria and/or their cristae. Five gene deficiencies caused MT-curling clearly demonstrating that mitochondrial dysfunction can affect MT bundles; in all cases, the phenotype was mediated by ROS, establishing oxidative stress as important mediator. Of these five genes, we discuss Sod2 as potential candidate that can explain MT-curling upon mitochondrial absence.

**Tab. 1.** Summary of findings reported here and by others about the employed genes, their functions, genetic tools and phenotypes. <u>Column 1</u>: the names of studied *Drosophila* genes; <u>column2</u>: their human orthologues; <u>column 3</u>: a crude description of the respective gene’s functions; <u>column 4</u>: the genetic manipulations employed (see abbreviations below; names of allele and knock-down lines provided in brackets); <u>column</u> <u>5</u>: summary of information from Fig.ZZ2 including the length of culture (DIV), the MDI trend (↓ decreased; ↑ increased; → no change), significance (asterisks) and value range from different LOF conditions (in brackets), as well as potential Trolox rescue as reported in Fig.ZZ5; <u>column 6</u>: summary of gene-linked neurological conditions or findings in mammalian models as described and referenced in the main text; <u>column 7</u>: summary of *in vivo* findings in *Drosophila* as described and referenced in the main text (findings obtained with the same tools as listed in column 4 are indicated by brackets); <u>column 8</u>: information relating to ROS upon gene manipulations in fly, mammalian models or humans as described and referenced in the Discussion. <u>Abbreviations:</u>; AD/R, autosomal dominant/recessive; CMT, Charcot Marie Tooth disease; DIV, days *in vitro*; EncP, encephalopathy; i-AAA proteases, <u>A</u>TPase <u>a</u>ssociated with various cellular <u>a</u>ctivities exposed to the inter-membrane space; IMM, inner mitochondrial membrane; ISC, iron-sulfur cluster; KD, knock-down (elav-Gal4-driven siRNA expression); KO, knock-out (complete or severe loss of function); ND, neurodegenerative disorder; NP, neuropathy; OA, optic atrophy; PD, Parkinson’s disease; PDC, pyruvate dehydrogenase complex; vs., versus; XD, X-chromosomal-linked dominant.

| gene | human orthologue | function | manipulation | MDI | neurological symptoms (OMIM ref. no.) | *in vivo* data in fly | effect on ETC or ROS |
| --- | --- | --- | --- | --- | --- | --- | --- |
| *Pdha1* | PDHA1 | catalytic component of PDC (pyruvate + CoA-SH → acetyl-CoA) | **KO (A); IR** (*HMC04032*) | 5 DIV ↑ **** (1.73-1.82)<br>Trolox rescue | neurological dysfunction (lactic acidosis/Leigh disease; XD#312170) | ↑CoA levels (*HMC04032*);<br>↓photoreceptors | <u>mouse/human</u>: increased ROS in cultured cells |
| *Ogdh1* | OGDH | Krebs cycle: α-ketoglutarate → succinyl-CoA | **KO** (*MI06026-TG4.1*), **KO/Df** | 5 DIV → (0.91-1.01) | movement disorders, ataxia, seizure (AR#203740) | embryonic lethal (*MI06026-TG4.1*); KD in photoreceptors: gradual loss of synaptic transmission | loss of OGDH: reduced O<sub>2</sub> consumption |
| *SdhA* | SDHA | Krebs cycle: succinate → fumarate; ETC: complex II component | **KO** (*1404*, *1110*) | 5 DIV ↑ **** (1.83-2.13)<br>Trolox rescue | ND with ataxia & OA (AD#619259) | gradual synapse loss (*1110* & *1404*); enhances ND phenotypes of *park* and *Sirup*/SDHAF4 | SDH complex deficiency causes ROS; synapse loss in fly is ROS-mediated; fly *SdhA* is upregulated by *Cnc/Nrf2* |
| *AtpSynC* | ATP5MC1/2/3 | F<sub>0</sub> components of ATP synthase; potential mPTP components | **KO** (*KG01914*); **KO/Df** | 1/3/5 DIV ↓→ (1.03-1.32) | ND: dystonia/SP (AD#619681) | larval lethal (*KG01914*) | <u>mouse</u>: loss of OXPHOS, protective against cellular stressors; <u>fly</u>: loss of cristae (*KG01914*); loss of OXPHOS |
| *mt:ATPase6* | MT-ATP6 |  | **KO** (*1*) | 5 DIV → (1.04) | severe neurological conditions (e.g. ataxia) | young flies: rounded cristae, ↓locomotion, seizure & lethality (all *1*) | <u>humans</u>: ROS overproduction (mPTP-related?); <u>fly</u>: loss of OXPHOS, glycolysis compensates (*1*) |
| *QIL1* | MICOS13 | key component required for MICOS complex formation | **IR** (*GLC01383*) | 5/6 DIV → (1.11) | ND with severe atrophies (#618329) | ↓QIL expression (>25%), aberrant mitochondria, ↑mitophagy, no cell death (all *GLC01383*) | reduced O<sub>2</sub> consumption; no reported ROS increase |
| *Opa1* | OPA1 | fusion of inner mito membrane; cristae formation | **KO** (*s3475*), **IR** (*HMS00349*) | 5 DIV/pre → (0.94-1.30) | NDs: Behr syndrome (AR#210000); OA (AD#165500; AD#125250) | ↓lifespan, but also rescue of tau-mediated ND (all *s3475*); ↑lifespan, ↑locomotion (*HMS00349*) | <u>mouse</u>: ↑ROS & ↑mtDNA damage, but also ↓ROS; <u>fly</u>: ↑ROS-mediated necrosis in eye (*s3475*); but also beneficial |
| *Marf* | MFN1/2 | fusion of outer mito membrane | **KO** (*B*), **IR** (*HMC03883*) | 5 DIV/pre → (0.60-1.05) | NDs: 2 CMTs (AD#609260; AR#617087); 2 NPs (AD#601152; AR#151800) | enhanced decline in HSP models, ↑ROS, ↑ER stress, ↑axon degeneration (*HMC03883*); but also improving frataxin, | <u>human</u>: ↑ROS; <u>mouse</u>: improved ROS tolerance, ↓ROS; <u>fly</u>: ↑ROS in nephrocytes (*HMC03883*), but also ameliorating |
|  |  |  |  |  |  | Huntington's & PD models, ↑lifespan, ↑locomotion | phenotypes in disease models |
| *Drp1* | DNM1L | mitochondrial fission | **KO** (T26) | 5 DIV/pre → (1.03-1.32) | NDs: EncP (AD/R#614388); OA (AD#610708) | ↓lifespan; but also ↑lifespan in proteasome-deficient model; rescue of ND in ALS model | <u>human/mouse</u>: ↓ROS |
| *YME1L* | YME1L1 | i-AAA protease (IMM) | **KO** (del) | 5 DIV ↓** (0.53) | ND: OA (AR#617302); late onset of ND in conditional mouse models | adults: mitos with aberrant cristae & aggregates, ↓locomotion, precocious death (all del) | <u>fly</u>: increased ROS at late stages; <u>yeast</u>: reduced OXPHOS; <u>human</u>: increased lactate/pyruvate levels |
| *sesB* | SLC25A4/5/6/31 aka ANT1-4 | ADP/ATP antiporter (IMM) | **IR1** (HMS01549), **IR2** (JF01528) | 4/5 DIV ↑**** (2.54-3.36) Trolox rescue | loose links to ND in humans; mouse myoblasts: ↓ETC dysfunction, ↓mito Ca<sup>2+</sup> response | ↓ATP, ↓mito Ca<sup>2+</sup> response, ↑autophagy, ↑ND | <u>mouse</u>: ↓glutathione, ↓mito peroxidases; <u>fly</u>: ↑H<sub>2</sub>O<sub>2</sub> |
| *Sod2* | SOD2 | Mn superoxide dismutase (superoxide → H<sub>2</sub>O<sub>2</sub>) | **KO** (n283) | 3 DIV ↑**** (2.78) Trolox rescue | loose links to ND in humans; early death & severe ND in mice with CNS-specific SOD2 loss | ↓life span, ↑ND/brain apoptosis, ↓locomotion (all n283); precocious axon decay, MT curling | <u>fly</u>: life span rescued by hypoxia; <u>mouse</u>: oxidative stress, proton leakage, DNA damage |
| *fh* | FXN | iron-sulfur-cluster formation | **KO** (1); **IR** (RNAi.A2) | 3/5/6 DIV ↑**** (2.90-2.93) Trolox rescue | Friedreich's ataxia (AR; 229300) | ↓photoreceptors, ↓ETC, ↓ATP, ↑Fe<sup>2+</sup> (all 1); ↓development/life span, ↓ETC (all RNAi.A2) | <u>fly</u>: ROS reports contradictory; <u>mouse</u>: ↓ISC, ↑iron, ↓Sod1/2, ↑Fenton react. |

## Results

### Loss of *PdhA* may cause MT-curling by impacting intra-mitochondrial metabolism

To assess the impact of mitochondrial physiology on axonal MT bundles, we selected candidate genes primarily based on their reported links to neurodegeneration and the availability of established genetic tools for their manipulation (see details below). We began our investigation with examples of factors that contribute to Krebs cycle function, starting with the pyruvate dehydrogenase complex (PDC) as the gate keeper which feeds the Krebs cycle with pyruvate derived from glycolysis or β-oxidation (Fig.1>*A-C/e*; letters behind the chevron indicate the grid coordinates provided in the figure). PDC is composed of multiples of three enzymes (E1-3) converting pyruvate and CoA-SH into acetyl-CoA and CO_2_ whilst generating NADH; PDHA (pyruvate dehydrogenase E1 subunit α1) is an obligatory subunit of the E1 enzyme which catalyses the first reaction step (Magistretti and Allaman, 2018; Naifeh et al., 2024; Patel et al., 2014).

In humans, PDHA1 mutations constitute ∼80% of cases of Leigh disease displaying with severe early-onset neurodegenerative symptoms, reduced PDC enzyme activity to ∼30%, and lactic acidosis with ∼4-fold increase in pyruvate and lactate plasma levels (Foucher and Tubben, 2024; Ganetzky et al., 2021; Gopal et al., 2023; Pavlu-Pereira et al., 2020). Rare patients with PDHB mutations show strong clinical overlap with PDHA mutant cases (Patel et al., 2012). In *Drosophila*, *in vivo* studies showed that Pdha1-deficient photoreceptor cells degenerate when challenged with light-induced activation (Jaiswal et al., 2015). Also strong knock-down of Pdhb (the β-subunit of E1) displayed clear neurodegenerative phenotypes (Dung et al., 2018).

To study loss of PDC function in primary neurons, we used homozygosis for the previously reported loss-of-function mutant allele *Pdha1^A^*(Liu et al., 2017; Yamamoto et al., 2014) and a knock-down line known to downregulate enzymatic activity of Pdha1 (Huang et al., 2022). In both cases, primary neurons showed a robust increase in MT-curling when analysed at 5DIV (Figs.2B, 3A), clearly confirming our hypothesis that axonal MT bundles can be affected by mitochondrial dysregulation. To test whether the phenotype of *Pdha1^A^* mutant primary neurons was ROS-dependent, we supplied the culture medium for the entire culture period with the ROS-scavenger Trolox (Chow et al., 1994). Trolox clearly suppressed the MT-curling phenotype (Fig.4C) suggesting harmful ROS dyshomeostasis to mediate the phenotype and confirming our prediction.

As will be discussed, reports of ROS involvement upon PDHA-deficiency in flies or mammals are inconsistent, but several studies find ROS production through intra-mitochondrial dysregulation (see Discussion and a summary of findings and reports in Tab.1). We would argue that any ROS leakage derived from such dysregulation can cause MT-curling only if mitochondria are present in axons, thus making PDC not a good candidate to explain axonal MT-curling when mitochondria stay in axons (see Introduction).

### Loss of Ogdh1 may downregulate the electron transfer chain rather than cause excessive ROS

Another Krebs cycle component we selected is the oxoglutarate dehydrogenase complex (OGDC) which converts α-ketoglutarate (aka oxoglutarate) to succinyl-CoA (Fig.1>*EF/fg*). Ogdh1/OGDH encodes the E1 component of the complex, which catalyses the initial step of the reaction (Nemeria et al., 2021).

In humans, OGDH deficiency causes movement disorders and hyperlactatemia (Yap et al., 2021b). In *Drosophila*, the *Ogdh1^MI06026-TG4.1^* loss-of-function allele is embryonic lethal, even if loss of Ogdh1 is restricted to the nervous system; its knock-down in photoreceptors causes progressive loss of synaptic transmission (Yap et al., 2021a; Yap et al., 2021b; Yoon et al., 2017; Tab.1). We generated *Drosophila* primary neurons (see Methods) from embryos carrying the *Ogdh1^MI06026-TG4.1^*allele in homozygosis or over deficiency and found no obvious increase in MT-curling at 5DIV, in one instance even a potential bias to reduce MT-curling (Fig.3B).

**Fig. 3.**
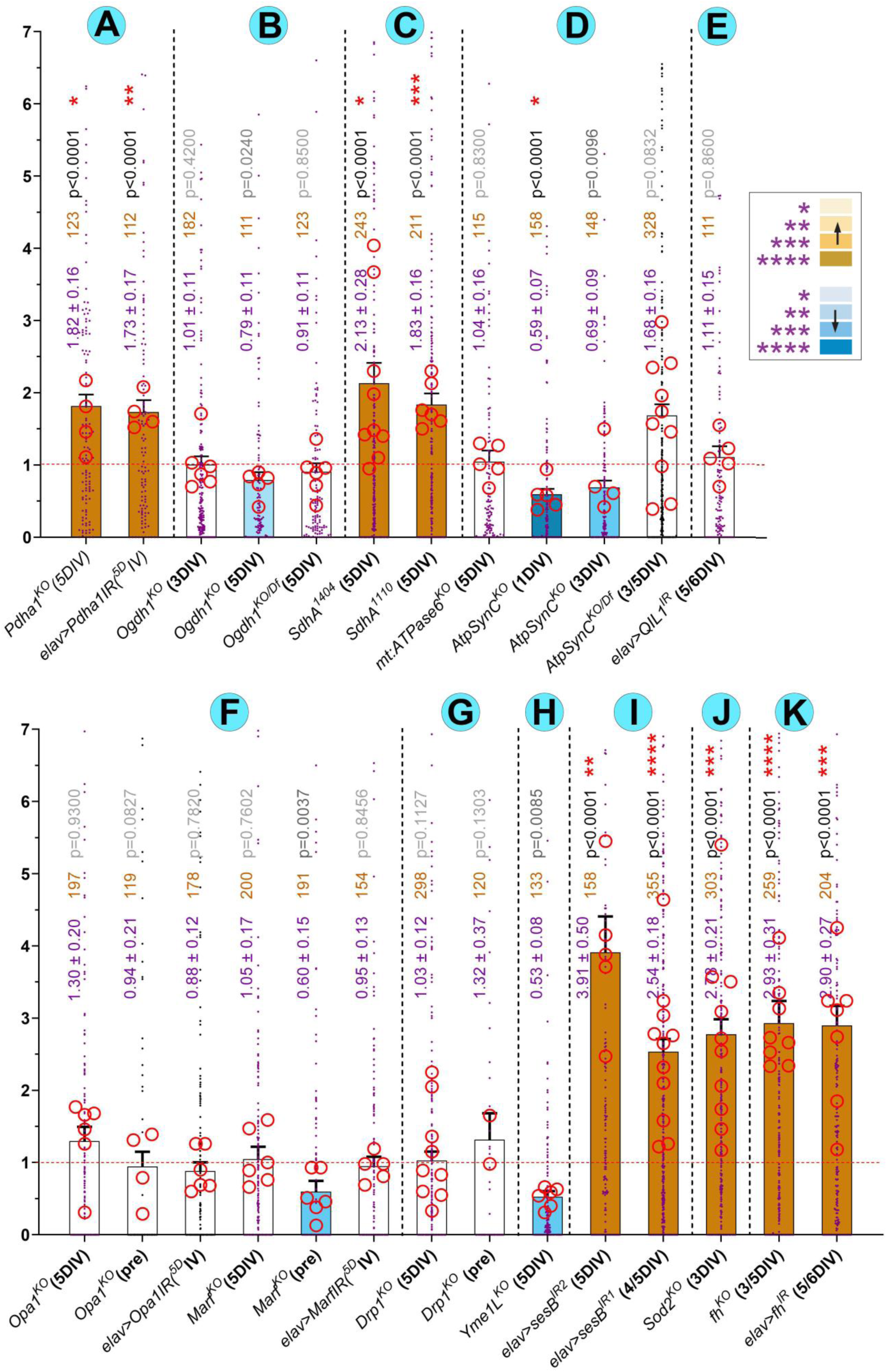
Quantification of MT-curling observed upon functional deficiency of factors involved in mitochondrial physiology. Bars indicate the mean ± SEM (numerical values shown in blue font) of MT-curling given as MT disorganisation index (MDI; the size of MT-curled areas relative to axon length); data are normalised to internal wild-type controls of each experiment (horizontal red dashed line); single data points are shown as blue dots and the means of replicates (independent coverslips from usually two to three independent replicates) are shown as red circles with their statistical significance (established using t-tests) indicated as red asterisks; numbers of assessed neurons are shown in orange, their p values relative to controls (established by Mann-Whitney tests) in black or grey; the grey-to-black intensity of statistical values and the brown/blue bar colour (see inset) reflect the respective degrees of significance relative to wild-type.

**Fig. 4.**
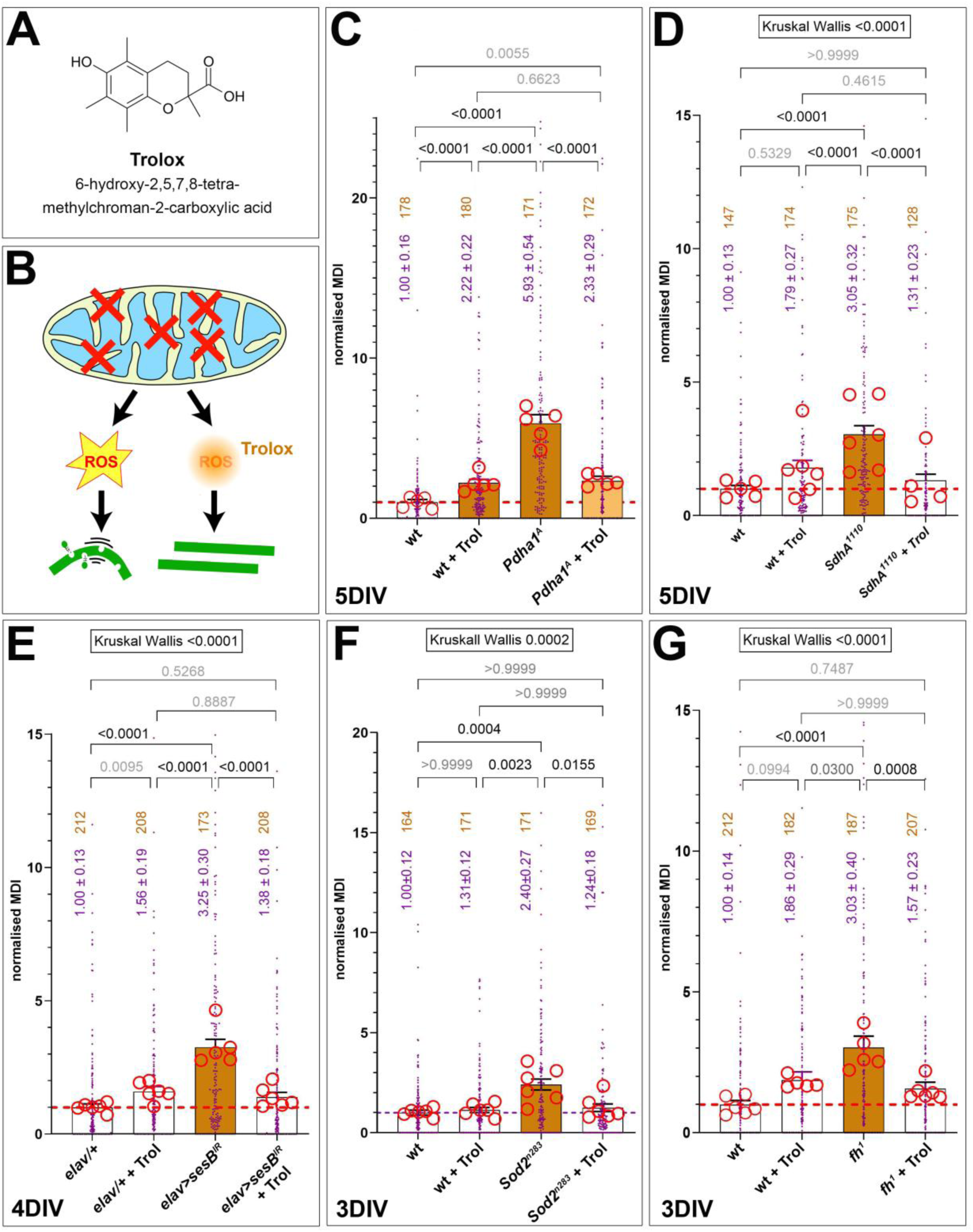
The phenotypes of all MT-curl-inducing conditions are rescued by Trolox. **A**) The chemical structure of Trolox. **B**) The design of Trolox experiments: mitochondrial phenotypes (red X) cause ROS (yellow star) which, in turn, causes curling and potential damage of MTs (left); upon application of Trolox (right), ROS is quenched and MT-curling abolished although the original mitochondrial phenotype is still present. **C-G**) Each graph represents one set of experiments composed of wild-type without Trolox (wt), wt with Trolox (+Trol) and the respective loss-of-function mutant condition (as indicated in C-G) without/with Trolox. Graphs are organised as explained in the legend of Fig.3. Kruskall-Wallis test results are indicated on top, Dunn’s multiple comparison results are indicated above bars; the intensity of brown fill-colour of bars reflects the degree of statistical significance relative to wild-type control.

As will be discussed, our finding seems to match well with reports about ROS in mammals or flies *in vivo*, especially with proposed roles of OGDH in ROS sensing and downregulation of the electron transfer chain (ETC) in times of oxidative stress (see Tab.1 and Discussion).

### SdhA deficiency causes MT-curling suppressed by Trolox

As the third Krebs cycle component we selected succinate dehydrogenase (SDH) which is an enzymatic complex formed by at least 4 distinct constituent subunits A to D. It participates in the Krebs cycle catalysing the step from succinate to fumarate. However, SDH is a special case in that it forms not only part of the Krebs cycle but also constitutes complex II of the ETC passing on electrons from its enzymatic reaction via ubiquinone to complex III (Fig.1>*EF/cd*; Al-Rasheed and Tarjan, 2018; Rutter et al., 2010).

In humans, SDHA mutations cause neurodegeneration, muscle weakness and tumour formation (Al-Rasheed and Tarjan, 2018; Hadrava Vanova et al., 2020; Patel et al., 2012; Rutter et al., 2010). The *Drosophila SdhA^1110^*and *SdhA^1404^* mutant alleles reduce SDH activity substantially causing recessive larval lethality; upon mosaic analysis, photoreceptor cells homozygous for these alleles showed gradual synapse loss (Mast et al., 2008). Furthermore, heterozygous deficiency of *SdhA* strongly enhances premature death of flies lacking the SDH assembly factor Sirup/SDHAF4 (Van Vranken et al., 2014) and *SdhA^1110^* genetically interacts with the Parkinson gene *park* enhancing its motility deficits (O’Hanlon et al., 2022).

Our analyses of primary neurons homozygous for the above-mentioned *SdhA^1404^* or *SdhA^1110^* mutant alleles, revealed a robust MT-curling phenotype at 5 DIV (Figs.2C, 3C). To test whether the phenotype of *SdhA*-deficient primary neurons was ROS-dependent, we applied the ROS-scavenger Trolox (Chow et al., 1994) which resulted in strong suppression of the MT phenotype (Fig.4D) suggesting oxidative stress as the mediator of MT-curling.

As will be discussed, the production of harmful ROS could be assembly defects of the SDH complex leaving iron-sulphur clusters of SDHB exposed (see Discussion).

### Loss of ATP synthase might dampen ROS abundance by downregulating ETC function

We next focussed on complex V of the ETC, also known as ATP synthase. ATP synthase is a multi-subunit complex clustered at the tips of cristae. Its channel-forming transmembrane sub-complex F_0_ permits the flow of protons across the inner membrane which provides the energy that drives ATPase activity of the enzymatic matrix-facing sub-complex F_1_ (Fig.1>*D/b*). APT synthase also plays important roles in cristae morphogenesis by mediating the extreme membrane curvature of their tips (Guo et al., 2017; Hahn et al., 2016; Zhou et al., 2015), and it is proposed to form a component of the mitochondrial permeability transition pore (mPTP; Bonora et al., 2022).

Here, we chose two key components of the F_0_ complex crucial for ATP synthase function: ATP5MC1-3 (ATPsynC in flies) constitutes the multimeric proton-conducting c-ring, whereas mitochondrially encoded MT-ATP6 (mt:ATPase6 in fly) closely associates with the c-ring and is required for its proton-conducting ability (Guo et al., 2017; Hahn et al., 2016; Zhou et al., 2015).

In humans, the dystonia-linked ATP5MC3^N106K^ mutation causes a reduction in ATP production and oxygen consumption (Neilson et al., 2022), and MT-ATP6 mutations are linked to severe neurological conditions (Dautant et al., 2018; Galber et al., 2021). In *Drosophila*, loss of ATPsynC (*ATPsynC^KG01914^*) causes a reduction in mitochondrial cristae and animals die as larvae after a prolonged developmental block (Lovero et al., 2018). Furthermore, ubiquitous expression of ATPsynC^N102K^ (mimicking human ATP5MC3^N106K^) caused strongly reduced ATP synthase activity coupled to lethality (Neilson et al., 2022). Individuals carrying the strong *mt:ATPase6^1^* loss-of-function allele (a G116E point mutation) were reported to be almost 100% mutant (homoplamic) leading to total loss of ATP synthase activity and aberrant mitochondria with severely rounded cristae; surprisingly, they can eclose but display a vastly reduced adult lifespan (Celotto et al., 2011; Celotto et al., 2006; Demine et al., 2019). We similarly observed that *mt:ATPase6^1^* mutant flies showed severely reduced mobility already at two weeks and a strong tendency to display seizures.

Our analyses of neurons homozygous for the above-mentioned null mutant allele *ATPsynC^KG01914^* (Lovero et al., 2018) revealed no obvious increase in MT-curling compared to wild-type controls at 3 or 5DIV; in some experiments it even caused a reduction in curling (Fig.3D). Likewise, *mt:ATP6^1^* mutant neurons failed to reveal any increases in MT-curling at 5 DIV (Fig.3D).

As will be discussed, our findings correlate well with the literature (Tab.1), although human MT-ATP6 mutations have late-onset pathology correlated with ROS production that might not be captured within the timeline of our experiments (see Discussion).

### Depletion of QIL1 or fission and fusion factors does not cause MT-curling within 5 days

We were surprised that the expected aberration of cristae upon loss of ATP synthase seems not to be a ROS-inducing condition. To challenge this finding, we studied further factors involved in cristae formation, namely the MICOS complex and Opa1. The MICOS complex is located at the neck of cristae (cristae junctions; Fig.1>*G/d*) required for their formation, the stabilisation of cristae junctions and the assembly of various protein complexes at this site (Mukherjee et al., 2021). When deleting the QIL1/MICOS13 subunit, the entire MICOS complex fails to form, causing reduced ETC activity and severe cristae aberrations (Mukherjee et al., 2021).

In humans, QIL1 mutations are linked to diseases with neurodegenerative traits (for example COXPD37; omim.org #<u>618329</u>). In *Drosophila*, knock-down of QIL1 in the nervous system and muscles reduced expression to under 25% accompanied by severe aberration of mitochondria, an increase in mitophagy, but no obvious induction of cell death (Guarani et al., 2015; Wang et al., 2020). We used the same knock-down construct in primary *Drosophila* neurons, but no obvious MT-curling phenotype was detectable at 5DIV (Fig.3E).

OPA1 is also positioned at the base of cristae and known to cause their disruption when dysfunctional (Fig.1>*G/c*; Quintana-Cabrera and Scorrano, 2023). However, OPA1 has an additional function in that it also regulates mitochondrial fusion (Fig.1>*D/a*). We therefore extended our study by including a second pro-fusion factor MFN (Fig.1>*B/a*) and the pro-fission factor DNM1L (Quintana-Cabrera and Scorrano, 2023).

In humans, all three pro-fission and -fusion factors have been linked to neurodegeneration (Tab.1; Chen et al., 2023a), and in *Drosophila* their losses cause lethality. However, studies of the human, mammalian or fly genes draw an inconclusive picture as to whether the pathologies involve harmful ROS production (Tab.1; see Discussion). We therefore assessed losses of Opa1 (*Opa1^s3475^* null allele and *elav>Opa1^IR^*), the MFN orthologue Marf (*Marf^B^* null allele and *elav>Marf^IR^*) and the DNM1L orthologue Drp1 (*Drp1^T26^*null allele) in *Drosophila* primary neurons. We observed no MT-curling at 5DIV, even when using pre-culture to exclude potential maternal rescue (see Methods; Figs.3F,G, 5B-D,F,G). Functional loss of the three factors in the assessed neurons was clearly indicated by fragmented mitochondria when depleting Opa1 or Marf (Fig.5B,C,F,G) and long stretches of continuous mitoTracker-labelled structures along primary axons Drp-deficient neurons (Fig.5D) as similarly reported for mammalian neurons (Berthet et al., 2014; Uo et al., 2009; Yu et al., 2011).

**Fig. 5.**
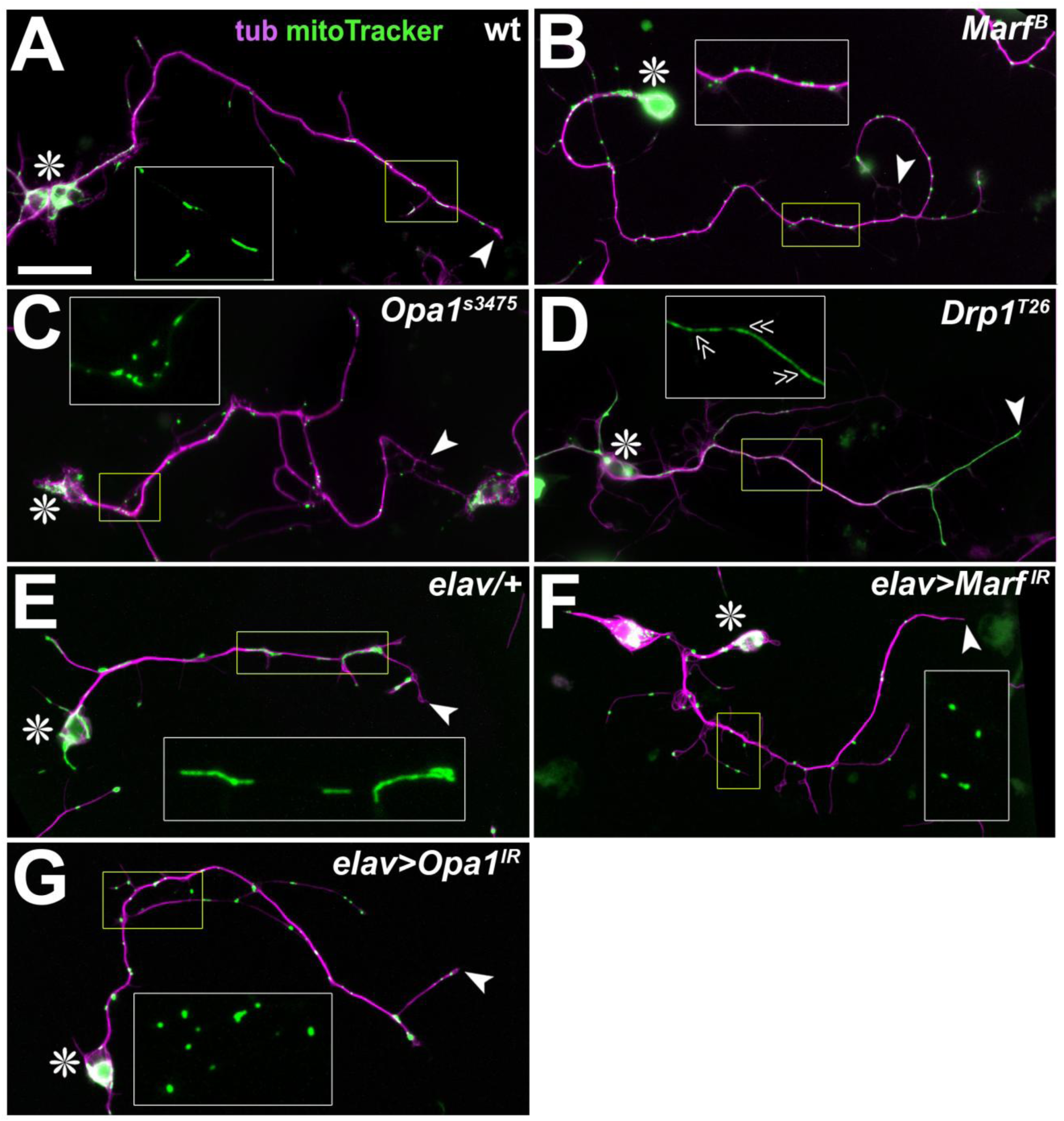
Genetic manipulations impairing mitochondrial fission/fusion processes. Neurons at 5 days *in vitro* (DIV) and stained with anti-tubulin (tub, magenta) and mitoTracker (green); they are wild-type (wt; A) or deficient for mitochondrial pre-fusion factors Marf (B,F) and Opa1 (C,G) and the mitochondrial pre-fission factor Drp1 (D); asterisks indicate cell bodies, arrow heads axon tips, yellow emboxed areas are shown as 2-fold enlarged insets (green channel only); the scale bar in A represents 30µm in A-D and 15µm in E-G; note that mitochondria tend to appear as dashed lines in controls (A), as sparse dots upon loss of fusion (B,C) and as a continuous structures excluded from many side branches upon loss of fission (‘double-chevrons’ in D). Quantification of MT-curling phenotypes under these mutant conditions are given in Fig.3.

Taken together, none of the factors involved in mitochondrial cristae formation, fission or fusion caused MT-curling in our standardised neuron system. As discussed, our data might therefore suggest that aberrant mitochondrial morphogenesis does not primarily lead to harmful ROS dyshomeostasis, at least over a period of 5 days.

### Loss of YME1L may reduce ROS production at early stages

The remaining factors addressed by our study cover a diverse range of further mitochondrial functions. For example, YME1L1 forms a homo-hexameric i-AAA protease which localises in the vicinity of the translocase complexes of the outer and inner mitochondrial membranes (TOMM and TIMM; Fig.1>*G/e*); YME1L1 plays important roles in proteolytic protein turn-over and the regulation of mitochondrial import and maturation (Kan et al., 2024).

Human patients homozygous for a hypomorphic YME1L1 mutation display onset of degeneration at childhood accompanied by an increase in lactate/pyruvate ratio indicative of glycolysis upregulation (Hartmann et al., 2016). In mice, ubiquitous knock-out is lethal due to heart dysfunction (Wai et al., 2015), and nervous system-specific deletion causes late-onset neurodegeneration and aberration of mitochondrial morphology and transport (Sprenger et al., 2019). Similarly, also *YME1L^del^* flies display neurodegeneration of photoreceptors, reduced locomotion and premature death at adult stage, correlating with severe mitochondrial pathology (cristae reduced and malformed; electron-dense inclusions due to unfolded protein stress) and an increase in ROS levels (Qi et al., 2016).

Surprisingly, we found that primary neurons homozygous for the protein null allele *YME1L^del^* displayed a consistent drastic reduction in MT-curling to about half of the values observed in parallel wild-type controls at 5DIV (Fig.3H). This might suggest a reduction in ROS production below base levels present in controls. As will be discussed, our findings align well with recent findings in yeast and with the late onset of *in vivo* pathologies in humans, mouse and fly (Tab1; see Discussion).

### Loss of non-redundant ANT in neurons causes MT curling

ANT (adenine nucleotide translocase) is a highly abundant mitochondrial protein in the inner mitochondrial membrane that acts as an ADP/ATP antiporter (‘SesB’ in Fig.1>*B/b*); it is also considered to be a component of the low-conductance mPTP helping to avoid mitochondrial calcium stress, and to mediate proton leakage involved in mitochondrial uncoupling (Bround et al., 2020; Chen et al., 2023b; Demine et al., 2019).

In humans, ANTs are discussed in the context of neurodegenerative diseases including AD and PD (Chen et al., 2023b). However, there are 4 functionally redundant ANT genes, of which the well-studied ANT1/SLC25A4 gene is expressed in brain, heart, muscles, lung and testis (linked to cardiomyopathy; OMIM #<u>103220</u>) and the poorly researched ANT3/SLC25A6 gene is ubiquitously expressed; very little is known about ANT2/SLC25A5 and ANT4/SLC25A31 (Chen et al., 2023b). In *Drosophila*, ANTs are encoded by the *sesB* gene (highly expressed in the nervous system) and the paralogous *Ant2* gene (mostly outside neural tissues; flybase.org, <u>FBgn0003360</u> and <u>FBgn0025111</u>; Li et al., 2022). Therefore, loss of SesB can be considered a total loss of ANT function in the nervous system; *in vivo* studies in *Drosophila* larvae or flies revealed decreased ATP production, reduced calcium response of mitochondria, elevated peroxide levels, clear signs of age-related neurodegeneration and enhanced autophagy (Celotto et al., 2006; DeVorkin et al., 2014; Terhzaz et al., 2010; Vartiainen et al., 2014).

In agreement with these *in vivo* findings, knock-down of *sesB* with two independent constructs showed a strong increase in MT-curling in primary neurons at 4 or 5DIV (Figs.2D, 3I). When treated with Trolox, the MT curling was reduced to control levels, suggesting harmful ROS dyshomeostasis as the curl-inducing factor (Fig.4E).

As will be discussed, the most likely cause of ROS production upon ANT deficiency is dysfunctional ETC in combination with the reduced ability of mitochondrial uncoupling and mPTP impairment, in both cases requiring mitochondria to be present in axons if it were to induce local MT-curling (see Discussion).

### Sod2 deficiency may explain MT-curling when mitochondria are absent

The mitochondrial manganese-containing superoxide dismutase SOD2 converts the highly toxic but short-lived superoxide (the main ETC-derived ROS; Andreyev et al., 2005; Murphy, 2008; Zorov et al., 2014) into the less aggressive H_2_O_2_ (Fig.1*>DE/e*) that is longer-lived and believed to diffuse into the cytoplasm contributing to local signalling (Fig.1*>A/f*; Palma et al., 2020).

Human SOD2 mutations are suggested to link to neurodegenerative diseases and conditions (Flynn and Melov, 2013; Houldsworth, 2024), although these associations are less clear than the ALS-links of SOD1 (Kim et al., 2020). SOD2 knock-out mice display increased apoptosis and urine acidity, cardiomyopathy and die after birth (Kokoszka et al., 2001; Li et al., 1995; Melov et al., 1998). If restricting SOD2 loss to the nervous system, severe neurodegeneration is observed (Melov et al., 1999; Oh et al., 2012; Qi et al., 2003). In the *Drosophila* brain, heterozygosity for *Sod2^Δ2^* and *Sod2^n64^* causes precocious axon decay and MT curling (Shields et al., 2025), and the *Sod2^n283^* allele caused reduced life span (rescued by hypoxia), early onset neurodegeneration at tissue and behavioural levels, enhanced apoptosis of brain cells, and sensitivity to oxidative stress (Dias-Santagata et al., 2007; Duttaroy et al., 2003; Paul et al., 2007; Piazza et al., 2009; Vrailas-Mortimer et al., 2011; Wicks et al., 2009).

We therefore used the *Sod2^n283^* mutant allele in homozygosis in primary neuron culture at 3DIV. These experiments revealed a strong increase in MT-curling, which was suppressed by Trolox, indicating involvement of harmful ROS (Figs.2E, 3J, 4F). Very similar results were recently reported by others using the *Sod2^Δ2^* and *Sod2^n64^* mutant alleles, clearly validating our findings (Shields et al., 2025).

As will be discussed, mitochondria lacking Sod2 function likely display enhanced electron leakage and harmful ROS production if present in axons. In contrast, mitochondria with normal Sod2 function may help to maintain cellular ROS homeostasis, and this role would be lacking and potentially cause MT-curling when mitochondria are absent from axons.

### Loss of frataxin may cause MT-curling by increasing ROS or via other mechanisms

FRATAXIN (FXN) is an iron-binding mitochondrial protein involved in the early steps of iron-sulphur cluster formation (ISCs; Fig.1*>C/b*; Monfort et al., 2022). ion-sulphur clusters are essential functional components of many proteins in mitochondria (including many ETC components), the cytoplasm and nucleus (Vallières et al., 2024). Although iron-sulphur cluster maturation and assembly into cytosolic and nuclear proteins takes place in the cytoplasm, the early steps always depend on mitochondria, making FXN a key factor in this functional context (Fan et al., 2022; Marelja et al., 2018; Marquez et al., 2023; Shi et al., 2021).

In humans, FXN mutations link to Friedreich’s ataxia as the most common form of autosomal recessive ataxia displaying with severe neurodegenerative pathology (Delatycki et al., 2000; Williams and De Jesus, 2024). In mouse models, this pathology is reproduced and correlates with iron accumulations and oxidative stress (Al-Mahdawi et al., 2006; Simon et al., 2004).

In *Drosophila*, Frataxin (Fh) loss causes strong neurodegeneration with dying-back symptoms of peripheral axons, aberrant mitochondrial appearance, enhanced mitophagy, increased iron uptake in mitochondria of the nervous system, and higher sensitivity to iron intake (Chen et al., 2016; Edenharter et al., 2018; Llorens et al., 2007; Navarro et al., 2015; Shidara and Hollenbeck, 2010). A reduced lifespan of flies was also observed when knocking down *frataxin* specifically in neurons (Anderson et al., 2005). From all these studies in flies, there are contradicting opinions as to whether Frataxin loss induces ROS (Marelja et al., 2018).

Employing genetic tools used for the above-mentioned *in vivo* experiments (Tab.1; lethal *fh^1^* S136R point mutation and *elav>fh^RNAi.A2^*) in primary neurons, we found a strong increase in MT-curling at 3 to 6DIV (Figs.2F, 3K). When applying Trolox, we found a clear reduction of the phenotype down to control levels (Fig.4G), indicating harmful ROS as the mediating factor.

As will be discussed, our data would indicate that loss of iron-sulphur clusters is a condition leading to harmful ROS. But mitochondria with functional Fh that remain in cell bodies would likely not cause a depletion of iron-sulphur cluster-containing proteins in the axonal cytoplasm because the transport distance in our cultured neurons are very short.

### Harmful ROS triggered by loss of Fh or SesB affects Eb1 amounts at MT plus ends

To test whether ROS produced upon loss of mitochondrial factors has an impact on other cell parameters, we used primary neurons mutant for *fh^1^* or with *elav-Gal4*-driven knock-down of *sesB* and assessed their morphological parameters. Neither axon length nor branch patterns (number of primary neurites) appeared affected (Fig.6). To assess whether other MT parameters were changed upon loss of these factors, we assessed the amount of Eb1 as an indicator of MT polymerisation (Hahn et al., 2021) and found a robust reduction in both cases. This reduction aligns with previous publications reporting reduced MT polymerisation upon ROS increase (Conze et al., 2025; Shields et al., 2025).

**Fig. 6.**
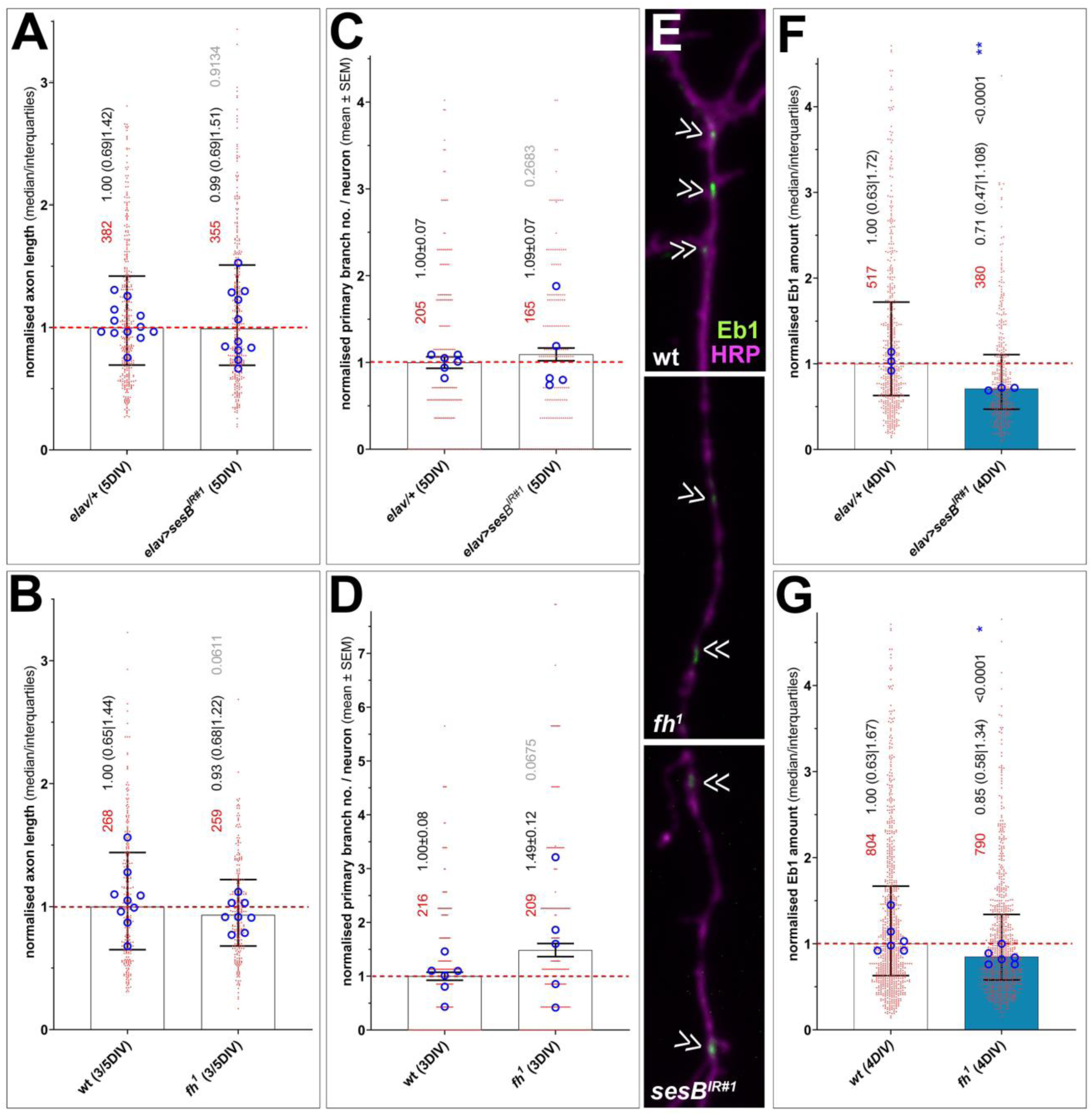
Functional losses of Fh and SesB do not affect neuronal morphology but trigger loss of Eb1 from polymerising MT tips. Quantification of axon growth (A,B), axon branching (number of primary neurites; C,D) and Eb1 amounts at MT plus ends (E-G) in neurons with loss of SesB (top) or Fh (bottom); images in E show axonal segments (stained for Eb1 in green and HRP in magenta); chevrons indicate Eb1 comets. In the graphs, bars indicate the median with quartile ranges or mean ± SEM (numerical values shown in black font); data are normalised to internal wild-type controls of each experiment (stippled line); single data points are shown as red dots (p values relative to controls established by Mann-Whitney tests are shown in black or grey) and the means of replicates (independent coverslips from usually two experimental repeats) are shown as blue circles with their statistical significance established using t-tests indicated as blue asterisks; numbers of assessed neurons are shown in red; the dark-blue bar colour reflects the degree of significance.

## Discussion

### Only certain forms of mitochondrial aberrations impact axonal MT bundles

Reports about mitochondrial physiology or dysfunction tend to be derived from highly diverse cellular models with very different metabolic footprints. These include liver cells with extreme metabolism rates or cancer cells which tend to lack oxidative phosphorylation. The execution of genetic manipulations in such diverse cell types often has inconsistent outcomes (see Discussion below and Tab.1); extrapolation of conclusions from one cellular context to another, and the integration of findings into conceptual understanding is therefore limited.

Here we used one consistent *Drosophila* primary neuron system to apply over 30 different genetic and experimental manipulations to study 13 genes important for mitochondrial physiology. We asked (1) which mitochondrial dysfunction can affect axonal MT bundles, (2) whether such effects are mediated by harmful ROS, and (3) through which mechanisms even the absence of mitochondria can cause MT-curling (see Introduction). Since MT bundles provide the lifelines of axons (see Introduction), potential pathological links from mitochondria to MTs might provide new mechanistic explanations for neurodegeneration. We chose the *Drosophila* primary neuron model because it has proven instrumental in the past when addressing other complex cell biological phenomena of axon biology, and could successfully demonstrate the fundamental principal that ROS impacts MT bundle organisation (Gonçalves-Pimentel et al., 2011; Hahn et al., 2021; Liew et al., 2025; Prokop et al., 2013; Qu et al., 2022; Shields et al., 2025; Voelzmann et al., 2024). Echoing this tradition, also our current approach delivered meaningful results:

Firstly, deficiencies of 5 out of 13 genes caused MT-curling, and all 5 conditions could be rescued by Trolox suggesting involvement of harmful ROS. Whilst being in line with the generally accepted view that certain mitochondrial aberrations can cause oxidative stress, they also clearly confirm that aberrant mitochondrial physiology impacts MT bundles with ROS being an important mediator. MT bundle deterioration provides therefore a potential downstream mechanism that links mitochondrial dysfunction to axon degeneration. The consistency of our findings might also suggest MT-curling as a complementary, easy to access readout for harmful ROS generation, although further validation including actual ROS measurements would be required.

Secondly, the results were surprising in that 8 out of the 13 gene deficiencies showed no obvious increases in MT-curling (Ogdh, QIL1, Opa1, Marf, Drp1, ATPsynC, mt:ATPase6, YME1L; Fig.3) despite their lethal or severely debilitating effects *in vivo*. It was even more surprising that, except Ogdh, all these factors are morphogenesis regulators of mitochondria and/or their cristae. Our finding suggests therefore that morphogenetic aberration is a condition that mitochondria can cope with, at least for several days. When mining the literature for these genes (see below; Tab.1), some of them are inconsistently discussed as to whether their pathological phenotypes involve ROS generation. But for all eight factors clear cases can be made in support of our findings - let alone the confidence provided by the unanimous and consistent nature of our own results. In the following we will discuss how our findings relate to the wider literature.

### Potential mechanisms that cause harmful ROS production upon loss of the 5 identified genes

We find ROS-induced MT-curling upon loss of Pdha1, although ROS seems not to feature in the current Pdha1-related *Drosophila* literature. However, ROS was reported from mammalian studies. Skin fibroblasts derived from PDHA-mutant patients showed increased mitochondrial but not cytoplasmic ROS levels (Glushakova et al., 2011), whereas increased cytoplasmic ROS levels were observed in PDHA1-deficient rat fibroblasts where Krebs cycle activity was partially maintained by glutamine-derived α-ketoglutarate (Fig.1>*A-E/g*; Wang et al., 2019). In studies of PDH-deficient mouse skeletal muscle, the ETC displayed low efficiency which was compensated by increased ETC activity - a potentially powerful constellation to cause electron leakage and generate ROS (Gopal et al., 2023). In our culture model, loss of Pdha1 might have even stronger impact on Krebs cycle activity and ETC because the loss of PdhA-derived pyruvate cannot be replenished from β-oxidation (Fig.1*>C/f*), since fatty acids are virtually absent from the culture medium (Else, 2020; Prokop et al., 2012; Schneider, 1964). Another potential mechanism for ROS generation could be the build-up of lactate (Fig.1*>A/e*) which causes harmful hyper-lactylation of proteins (Chapp et al., 2021; Wang et al., 2024; Yang et al., 2024). However, build-up of lactate is highly unlikely in *Drosophila* primary neurons which are grown in lactate-free culture medium (Schneider, 1964), leaving impacts of Krebs cycle aberration on the ETC as the more likely option.

SdhA-linked pathology in *Drosophila* has been linked to ROS *in vivo*, which agrees with our findings. For example, antioxidants could reduce synapse loss in *SdhA^1110^* and *SdhA^1404^* mutant photoreceptors (Mast et al., 2008), and SdhA is part of the Cnc/Nrf2-mediated oxidative stress response pathway (Tsakiri et al., 2019a). For vertebrates, we found reports of ROS elevation so far only for loss of subunits B, C and D (Goffrini et al., 2009; Guzy et al., 2008; Hadrava Vanova et al., 2020; Ishii et al., 2011; Li et al., 2019; Owens et al., 2012), but there were also arguments against ROS. For example, SDHB-deficient chromaffin cells had reduced oxygen consumption (Kľučková et al., 2020), SDH was suggested to act as a ROS sensor dampening Krebs cycle and ETC activity upon elevated H_2_O_2_ levels (Nulton-Persson et al., 2003; Nulton-Persson and Szweda, 2001). Furthermore, complex II is often missing from respirasome super-complexes where complex I can directly reduce ubiquinone (‘Q’ in Fig.1>*EF/cd;* Enriquez and Lenaz, 2014; Hadrava Vanova et al., 2020; Waltz et al., 2024). Regarding the mechanisms of ROS production, Krebs cycle aberration is not a very likely cause since there are various compensatory pathways. For example, SDHB loss in mouse chromaffin cells causes depletion of malate (Fig.1>*D/d*), but Krebs cycle activity is upheld by pyruvate carboxylase-derived oxaloacetate (‘Pcb’ in Fig.1>*C/de*; Lussey-Lepoutre et al., 2015). The malate-aspartate shuttle (Fig.1>*B/cd*; Broeks et al., 2021; Koch et al., 2024), which seems present in *Drosophila* (Curcio et al., 2020; Karp et al., 2017), could even ensure malate dehydrogenase-mediated NADH production (‘Mdh2’ in Fig.1>*D/d*). In our view, an attractive explanation for ROS production upon loss of SDHA is the suggestion that SDH complex formation is impaired in ways that leave the iron-sulphur clusters of SDHB exposed, thus providing a potential ROS-producing source (Hadrava Vanova et al., 2020; Lemarie et al., 2011).

Our results for SesB align with *in vivo* reports for *Drosophila* that its loss causes elevated peroxide levels (Celotto et al., 2006; DeVorkin et al., 2014; Terhzaz et al., 2010; Vartiainen et al., 2014). This is even clearer from mammalian studies. For example, ANT1/2-deficient mouse myoblasts displayed ETC dysfunction as well as reduced glutathione levels and mitochondrial peroxidase activities (Flierl et al., 2022). Knock-down of ANT2 in MCF-7 cells caused significant increase in ROS (Kretova et al., 2014), isolated mitochondria from ANT1-deficient mice had significantly increased hydrogen peroxide production (Esposito et al., 1999), ANT1 overexpression in rat heart protects from ROS damage (Klumpe et al., 2016), and ETC components and activities are upregulated in ANT1/2 double-mutant mouse liver (Kokoszka et al., 2004). This said, the ETC was downregulated in ANT1-deficient mouse muscle fibres (Graham et al., 1997) and ANT1 loss was even suggested to be beneficial for cell stresses (Lee et al., 2009), but these studies did not address the potential redundancy of ANT genes. In our view, the most likely cause of ROS production is the dysfunctional ETC in combination with the reduced ability of mitochondrial uncoupling and mPTP impairment.

Our results for Sod2 appear the easiest to explain and align well with very recent data generated in the same *Drosophila* primary neuron system (Shields et al., 2025). Mitochondria of mice lacking SOD2 display reduced respiration, a sensitised transition pore, increased proton leakage, as well as a reduced expression of Krebs cycle and ETC enzymes (including Sdh and aconitase; ‘Aco’ in Fig.1); knock-out mice display, oxidative stress and genomic DNA damage (Kokoszka et al., 2001; Li et al., 1995; Melov et al., 1998). Increased cytoplasmic ROS levels upon SOD2 loss are likely the consequence of risen intra-mitochondrial superoxide which fails to convert to H_2_O_2_ and reacts with iron-sulphur clusters (Fig.1*>CD/c*), thus damaging ETC components and enhancing electron leakage (Palma et al., 2020). Further impact may derive from SOD2-mediated production of H_2_O_2_ which is far more diffusive and long-lived than superoxide and a known signalling molecule diffusing to the surrounding cytoplasm; since SOD2 activity is regulated by the metabolic and redox state of mitochondria it could act as an integrating sensor, and its H_2_O_2_-mediated signalling might help to maintain ROS homeostasis in the surrounding cytoplasm (Palma et al., 2020). Importantly, this signalling role would be affected if mitochondria are absent from axons, potentially explaining why failed mitochondrial transport causes MT-curling (Liew et al., 2025).

As mentioned before, the potential involvement of ROS in *fh*-linked phenotypes in *Drosophila* is controversially reported and debated (Marelja et al., 2018). Various mechanisms could explain potential ROS increase upon loss of Frataxin. Firstly, the failure of proper iron-sulphur cluster formation will impact on many mitochondrial proteins especially of the ETC which might cause increased electron leakage, hence ROS production. Secondly, Frataxin-deficient mitochondria displayed increased iron levels (Chen et al., 2016; Navarro et al., 2015) which, in turn, inactivated Sod2 with the respective knock-on effects on MTs (Marelja et al., 2018). High iron levels also trigger ferrotopsis-related ROS-inducing processes including the Fenton reaction which generates highly reactive hydroxyl radicals, also observed in Friedreich’s ataxia (Anderson et al., 2008; Costa et al., 2023; La Rosa et al., 2021).

Taken together, the losses of all 5 factors likely cause harmful ROS through intra-mitochondrial dysfunction, meaning that mitochondria must be present in axons to cause cytoplasmic ROS that can trigger MT-curling. However, the role of Sod2 in generating H_2_O_2_ as a function of mitochondrial physiology might establish mitochondria as sensors and regulators of local cytoplasmic ROS homeostasis; this might explain the phenomenon that even absence of mitochondria can cause MT-curling (Liew et al., 2025).

### Non-curl-inducing conditions

Eight out of 13 gene deficiencies failed to induce MT-curling. Functional loss of ATPsynC or YME1L even showed signs of reduced MT-curling (Fig.3G,H), suggesting a potential drop in ROS levels below baseline. Because *Drosophila* primary neurons are grown in an environment with higher oxygen levels than experienced *in vivo*, their baseline levels of ROS might be slightly elevated, thus setting a bias for MT-curling that can be reduced.

In agreement with our findings for Ogdh1, also OGDHL-deficient human neuroblastoma cells were shown to reduce oxygen consumption (Yap et al., 2021a). OGDH loss affects the Krebs cycle with downstream effects like causing mTORC1 activation (Yoon et al., 2017), although cycle activity can be partly maintained: for example via import of malate (Fig.1>*AB/c*; Allen et al., 2016; Yoon et al., 2017) or via methionine catabolism to generate succinyl-CoA (not shown; Yap et al., 2021a). Notably, OGDH was shown to be inactivated through glutathionylation in response to heightened H_2_O_2_ levels, thus acting as a ROS sensor that can down-regulate NADH production and OXPHOS (Applegate et al., 2008; Nulton-Persson et al., 2003; Nulton-Persson and Szweda, 2001). Loss of Ogdh1 might mimic this silencing effect, expected to cause a reduction or at least no increase in ROS production, as seen in our studies.

Consistent with the lack of MT-curling we observed upon QIL1 knock-down, also MICOS13-deficient cells were reported to display reduced oxygen consumption (Guarani et al., 2015; Kishita et al., 2020) and no studies seem to suggest ROS involvement (Gödiker et al., 2018; Guarani et al., 2016; Russell et al., 2019; Zeharia et al., 2016).

Matters are less clear for the fission and fusion factors. In *Drosophila*, Drp1 loss decreases lifespan (Rana et al., 2017), but there seem to be no reports of ROS-induced neurodegeneration; instead Drp1 deficiency was shown to rescue longevity in a proteasome-deficient model (Tsakiri et al., 2019b), and dominant-negative Drp1 was beneficial in ALS models (Altanbyek et al., 2016). The *Opa1^s3475^* mutant allele was shown to cause elevated ROS, shorter lifespan and necrosis of support cells in the fly eye (Tang et al., 2009; Yarosh et al., 2008), and Opa1 knock-down caused axon degeneration (Cao et al., 2017). In other reports, *Opa1^s3475^*reduced toxicity in a *Drosophila* Alzheimer model (DuBoff et al., 2012), and *Opa1* knock-down in muscles increased lifespan and improved locomotor activity (Tapia et al., 2021). Similarly, functional loss of Marf causes precocious axon degeneration (Cao et al., 2017), promotes decline in spastic paraplegia models (Fowler and O’Sullivan, 2016), triggers ROS-increase in nephrocytes (Zhu et al., 2024) and leads to ER stress and fragmentation (Debattisti et al., 2014). But it was also shown to rescue frataxin-induced glial degeneration (Edenharter et al., 2018), alleviate ROS-mediated rhabdomere degeneration in Huntington’s disease models (Campesan et al., 2023), reduce pink/parkin-induced ER stress in a fly Parkinson model (Celardo et al., 2016; but also see Basso et al., 2018) and increase lifespan and locomotor activity (Rana et al., 2017; Tapia et al., 2021). Also in mammalian studies, some report ROS increase upon mitochondrial fragmentation caused by OPA1 or MFN loss (Ježek et al., 2018; Millet et al., 2016; Zhang et al., 2017), whereas others report that DNM1L loss causes proton leakage and a reduction in ROS (Galloway et al., 2012; Kolac et al., 2023), that OPA1-deficient mouse embryonic fibroblasts and hepatocytes have reduced ROS (Lee et al., 2023; Liang et al., 2024), that MFN1-deficient myocytes display normal mitochondrial physiology and improved ROS tolerance (Papanicolaou et al., 2012), and that loss of MFN2 in macrophages have decreased ROS production (Tur et al., 2020). Our experiments for all these factors consistently show lack of MT-curling arguing against ROS increase. We propose that local maintenance mechanisms including mitochondrial protease systems and mitochondria-derived vesicles might be able to keep mitochondrial physiology in balance for an extended period (Misgeld and Schwarz, 2017).

Lack of MT-curling upon functional deficiency of ATP synthase aligns with findings in the literature. For example, blocking mouse ATP synthase affects respirasome assembly and metabolically protects neurons against cellular stresses (Formentini et al., 2014; García-Aguilar and Cuezva, 2018), and cells carrying the dystonia-linked ATP5MC3^N106K^ mutation display reduced ATP production and oxygen consumption (Neilson et al., 2022). The surprising survival of *mt:ATPase6^1^* mutant individuals into adult flies was explained by mitochondrial uncoupling (Demine et al., 2019) and glycolytic ATP production (Fig.1>*A/de*; Celotto et al., 2011), none of which would suggest ROS production. However, ROS overproduction has been reported for neurodegeneration-linked human MT-ATP6 mutations (Dautant et al., 2018; Galber et al., 2021), in one case involving overproduction of SOD1 and 2 (Geromel et al., 2001). Such effects might relate to roles of the ATP synthase in forming the high-conductance mitochondrial permeability transition pore (mPTP; Bonora et al., 2022) causing late-onset effects not covered by our experimental schedule.

As explained in the Results section, also loss of Yme1L causes late onset of ROS and neurodegeneration in fly and mouse alike, whereas we find that loss of Yme1L causes a strong reduction of MT-curling at 5 DIV. Our results are perhaps best explained by findings in yeast, where loss of *Yme1* causes a severe drop in levels of various ETC components expected to reduce activities of complexes II, III and IV (Kan et al., 2022) and, hence, reduce ROS production. We propose therefore that the loss of Yme1 protease activity causes reduced processing of ETC components leading to early ROS reduction, whereas its roles in mitochondrial quality control, i.e. to remove protein aggregates, become more relevant over a longer time period, potentially masked by other maintenance mechanisms including the shedding of mitochondria-derived vesicles (Held and Houtkooper, 2015; Mattedi et al., 2023; Misgeld and Schwarz, 2017), thus explaining late-onset ROS not seen in our cultures.

### Conclusions and future directions

The approach taken here was clearly able to answer the posed questions regarding the ROS-mediated impact of mitochondrial dysfunction on MT bundles. This suggests axonal MT bundles as potential downstream targets in mitochondrial pathology, that would provide a mechanism for axon degeneration. Our data might also suggest Sod2 as a potential candidate explaining MT bundle deterioration upon mitochondrial absence. Further thought-provoking observations were made, such as the absence of MT-curling when affecting mitochondrial morphogenesis, or the strong reduction of curling upon loss of QIL1. Importantly, the *Drosophila* primary neuron system is well-suited to validate findings and test deduced hypotheses efficiently, for example using double- or triple-mutant conditions to clarify compensatory metabolic pathways. Furthermore, findings can be easily validated *in vivo* using the same genetic tools as in culture. This provides promising means to refine our understanding of mitochondria in axons.

## Methods

### Fly lines

The wild-type control used throughout the project was the *Drosophila melanogaster* Oregan R strain. Loss-of-function mutant strains were (in alphabetical order): ***ATPsynC^KO^*** (*ATPsynC^KG01914^*; P-element insertion in non-coding 5’ exon generating a protein null; BDSC#13923; Lovero et al., 2018); ***ATPsynC^Df^*** (*Df(3R)Exel6218*; uncovering ATPsynC; BDSC#7696; Parks et al., 2004); ***ATP6^KO^*** (*mt:ATPase6^1^*; lethal G116E point mutation; BDSC#95253; Celotto et al., 2006); ***Drp1^KO^***(*Drp1^T26^*; lethal allele; BDSC#3662; Verstreken et al., 2005); ***fh^KO^*** (*fh^1^*; lethal S136R point mutation; BDSC#67161; Chen et al., 2016); ***Marf^KO^*** (*Marf^B^*; amorphic allele; BDSC#67154; Sandoval et al., 2014); ***Ogdh1^Df^*** (*Df(3L)Exel7253*; BDSC#7938; Ryder et al., 2007); ***Odgh1^KO^*** (*Mi{Trojan-GAL4.1}Ogdh1^MI06026-TG4.1^* aka *dOgdh-T2A-Gal4*; BDSC#77497; Yoon et al., 2017); ***Opa1^KO^*** (*Opa1^s3475^*aka *Opa1^ex2^*; BL #12188; strong loss of function due to P-element insertion; Spradling et al., 1999; Yarosh et al., 2008); ***Pdha1^KO^*** (*Pdha1^A^* aka *l(1)G0334A*; lethal G126E point mutation; BDSC#52370; Yamamoto et al., 2014); ***SdhA^1404^*** (lethal V445E point mutation; BDSC#81120; Mast et al., 2008); ***SdhA^1110^*** (lethal E288K point mutation; BDSC#51659; Mast et al., 2008); ***Sod2^KO^*** (*Sod2^n283^*; BL#34060; 167bp deletion removing part of the first exon and intron; Duttaroy et al., 2003); ***YME1L^KO^*** (*YME1L^del^*; 2kb deletion removing most of the coding region; BDSC#95273; Qi et al., 2016). Knock-down experiments were performed using the Gal4/UAS system (Elliott and Brand, 2008) employing the second-chromosomal ***elav-Gal4*** driver (BDSC #8765) in combination with the following transgenic UAS constructs: ***fh^IR^*** (*P{UAS-fh.RNAi.A}2*; BDSC#24620; Anderson et al., 2005); ***Marf^IR^*** (*HMC03883*; BL#55189; Perkins et al., 2015); ***Opa1^IR^*** (*HMS00349*; BDSC#32358; Perkins et al., 2015); ***Pdha^IR^*** (*P{TRiP.HMC04032}*; BDSC#55345; Perkins et al., 2015); ***QIL1^IR^*** (*P{TRiP.GLC01383}*; expression reduced to about 25%; loss of cristae junctions; BDSC#44364; Guarani et al., 2015; Perkins et al., 2015); ***sesB^IR1^*** (*P{TRiP.HMS01549};* BDSC#36661; Perkins et al., 2015); ***sesB^IR2^*** (*P{TRiP.JF01528};* BDSC#31077; Perkins et al., 2015). Green balancer chromosomes used to identify mutant or construct-expressing embryos were readily available *FM7*, *CyO or TM3* balancers carrying *twi-Gal4 or Kr-Gal4* in combination with *UAS-GFP* or carrying a *Df-GFP* fusion construct (Casso et al., 2000; Halfon et al., 2002; Le et al., 2006).

### *Drosophila* primary cell culture

*Drosophila* primary neuron cultures were performed as published previously (Prokop et al., 2012; Voelzmann and Sánchez-Soriano, 2022). In brief, stage 11 embryos were treated for 1 min with bleach to remove the chorion, sterilized for ∼30 s in 70% ethanol, washed in sterile Schneider’s/FCS, and eventually homogenized with micro-pestles in 1.5 centrifuge tubes containing 21 embryos per 100μl dispersion medium and left to incubate for 5 min at 37°C. Cells were washed with Schneider’s medium (Gibco), spun down for 4 mins at 650g, supernatant was removed and cells re-suspended in 90µl of Schneider’s medium containing 20% fetal calf serum (Gibco). 30μl drops were placed on cover slips. Cells were allowed to adhere for ∼2hrs either directly on glass or on cover slips coated with a 5 µg/ml solution of concanavalin A and then grown as a hanging drop culture for hours or days at 26°C as indicated in each experiment. To abolish maternal product deposited by heterozygous mothers in their oocytes (Prokop, 2013), we used a pre-culture strategy (Prokop et al., 2012; Sánchez-Soriano et al., 2010) where cells were kept for 5 days in a tube before they were plated on a coverslip [indicated as ‘(pre)’ in Fig.3].

Cells were treated with 100 µM Trolox (Sigma; stepwise diluted from a 100mM stock solution in ethanol; controls were cultured in medium with the equivalent amount of ethanol). In later experiments, Trolox was directly added to the medium and then sterile-filtered. 100 µM diethyl-maleate (DEM) was directly added to the medium and then sterile-filtered. Either agent was kept on the cells throughout the entire period.

### Staining procedures

Cultured neurons on coverslip were fixed for 30 minutes using a drop of 4% paraformaldehyde (PFA) and 0.05% glutardialdehyde in 0.05M phosphate buffer (PB; pH 7–7.2). Cells were washed with 0.5% Tergitol solution in 0.05M PB for 20 minutes (one exchange). The cells were incubated for 2 hrs in a 200µl drop of anti-tubulin primary antibodies diluted 1:500 in PB (clone DM1A, mouse, Sigma or clone YOL1/34, rat, Antibodies.com). Cells were washed with PBS and then incubated for 1.5 hrs with secondary FITC- or Cy3-conjugated anti-mouse antibodies (donkey, Jackson ImmunoResearch, 1:200 in PBS). To image cell morphology and identify neurons, cells were co-labelled with TRITC/Alexa647-, FITC- or Atto647N-conjugated anti-HRP (goat, Jackson Immuno Research, 1:100). For Eb1 comet analysis, cells were fixed for 10 min in the freezer with pre-cooled Plus-Tip Fixative (90% methanol, 3% formaldehyde, 5 mM sodium carbonate, pH 9), then washed with PBS and stained as described above with anti-DmEb1 (gift from H. Ohkura; rabbit, 1:500; Elliott et al., 2005). For the visualisation of mitochondria, cell cultures were incubated with 1µM MitoTracker Red CMXRos (Invitrogen; Klionsky et al., 2012) for 30min at room temperature (RT); stock solutions were prepared in DMSO and diluted in cell culture medium to the final concentration. Following incubation, cultures were then fixed and stained following the procedures below. Specimens were embedded in ProLong Gold Antifade mounting medium (Invitrogen) on microscope slides. The embedded slides were left to dry overnight in the dark before imaging.

### Imaging and data analysis

Neurons were visualised using a compound fluorescence microscope (BX50WI or BX51; Olympus) and images of single neurons were captured using nijiBlueBox and the MatrixVision mvBlueFox3-M2 2124G camera at 100x magnification. Images were analysed using the FIJI/ImageJ software: to determine the degree of MT disorganisation in axons we used the “MT disorganisation index” (MDI; Qu et al., 2017): the axon length (from cell body to tip of the most distant microtubule) was measured using the segmented line tool; area of disorganisation was measured using the freehand selection tool; this value was then divided by the the product of axon length multiplied with 0.5 μm (arbitrary axon diameter, thus approximating the expected area of the axon if it were not disorganised); for axon branching data, primary neurites containing a microtubule core of at least 10 µm and branching off the longest neurite were counted. For the Eb1 comet analysis, length and mean intensity of the Eb1 comets was measured using the line tool in FIJI; Eb1 amount was calculated by multiplying comet length with mean intensity (Hahn et al., 2021). All data were normalised against their respective controls. In each experiment, usually three slides per genotype were analysed aiming to image ∼30 isolated neurons per slide. Experiments were repeated at least once, data pooled. MDI data were usually not normally distributed but nevertheless plotted as mean ± SEM to avoid misleading median values of zero. Most experiments had only two groups and were assessed using Mann–Whitney Rank Sum tests, experiments with more then two group using Kruskal–Wallis one-way ANOVA with *post hoc* Dunn’s test. Means of single slides were used to generate super-plots (Lord et al., 2020) and assessed using standard t-tests. The data used for our analyses will be made available on request.

## Abbreviated gene names

Acon: mitochondrial Aconitase
ABCB7: ATP binding cassette subfamily B member 7 Aralar, mitochondrial aspartate/glutamate carrier
ATP6: mitochondrial ATPase subunit 6 (*Drosophila* name: mt:ATPase6)
ATPsynC: ATP synthase, subunit C
DIC: Dicarboxylate carrier 1
F_0/1_: F_0_ and F_1_ subunits of the ATP synthase
Fh: Frataxin
Gdh: Glutamate dehydrogenase
Gls: Glutaminase
Got2: Glutamate oxaloacetate transaminase 2
HRP: horseradish peroxidase
i-AAA: <u>A</u>TPase <u>a</u>ssociated with various cellular <u>a</u>ctivities exposed to the inter-membrane space
ISC: ion-sulfur cluster
Ldh: Lactate dehydrogenase
Marf: Mitochondrial assembly regulatory factor
MIA40: Mitochondrial intermembrane space Import and Assembly 40
MICOS: mitochondrial contact site and cristae organizing system
Mpc1: Mitochondrial pyruvate carrier
OGDC: Oxoglutarate dehydrogenase complex
Ogdh1: Oxoglutarate dehydrogenase 1
Opa1: Optic atrophy 1
QIL1: subunit of MICOS, orthologue of MICOS13
Pcb: Pyruvate carboxylase
PDC: Pyruvate decarboxylase complex
PdhA: Pyruvate dehydrogenase E1 alpha subunit 1
PerX: peroxisomes
SesB: stress-sensitive B; orthologue of ANT (SLC25A4-6)
Sod2: Superoxide dismutase 2 (Mn)
TIM22/23: Mitochondrial import inner membrane translocase subunits 22/23
TOM40: translocase of the outer mitochondrial membrane 40
YME1L: YME1 like ATPase; i-AAA protease of the inner mitochondrial membrane

## Acknowledgements

This work was made possible through support by the BBSRC to A.P. (BB/P020151/1) and parent support to Y.-T.L.. The Fly Facility has been supported by funds from The University of Manchester (https://www.bmh.manchester.ac.uk/research/support/funding/strategic) and the Wellcome Trust (087742/Z/08/Z; AP). Stocks obtained from the Bloomington Drosophila Stock Center (NIH P40OD018537) were used in this study. The funders had no role in study design, data collection and analysis, decision to publish, or preparation of the manuscript.

